# Elastase-2 knockout mice display anxiogenic‐ and antidepressant-like phenotype: putative role for BDNF metabolism in prefrontal cortex

**DOI:** 10.1101/187831

**Authors:** CRAF Diniz, C Becari, A Lesnikova, C Biojone, MCO Salgado, HC Salgado, LBM Resstel, FS Guimaraes, E Castren, PC Casarotto, SRL Joca

**Affiliations:** Department of Pharmacology, Ribeirão Preto Medical School, University of Sao Paulo, Ribeirão Preto, SP, Brazil.; Department of Physiology, Ribeirão Preto Medical School, University of Sao Paulo, Ribeirão Preto, SP, Brazil.; Department of Physics and Chemistry, School of Pharmaceutical Sciences of Ribeirão Preto, University of Sao Paulo, Ribeirão Preto, SP, Brazil.; Department of Cardiovascular Diseases, Mayo Clinic, Rochester, MN, USA.; Neuroscience Center, Helsinki Institute of Life Science, University of Helsinki, Helsinki, Finland.; Translational Neuropsychiatric Unit, Department of Clinical Medicine, Aarhus University, Denmark

## Abstract

Several pieces of evidence indicate that elastase-2 (ELA2; chymotrypsin-like ELA2) is an alternative pathway to the generation of angiotensin II (ANG II). Elastase-2 knockout mice (ELA2KO) exhibit alterations in the arterial blood pressure and heart rate. However, there is no data on the behavioral consequences of ELA2 deletion. In this study we addressed this question, submitting ELA2KO and wild-type (WT) mice to several models sensitive to anxiety‐ and depression-like, memory, and repetitive behaviors. Our data indicates a higher incidence of barbering behavior in ELA2KO compared to WT, as well as an anxiogenic phenotype, evaluated in the elevated plus maze (EPM). While a decrease in locomotor activity was observed in ELA2KO in EPM, this feature was not the main source of variation in the other parameters analyzed. The marble burying test (MBT) indicated increase in repetitive behavior, observed by a higher number of buried marbles. The actimeter test indicated a decrease in total activity and confirmed the increase in repetitive behavior. The spatial memory was tested by repeated exposure to the actimeter in a 24h interval. Both ELA2KO and WT exhibited decreased activity compared to the first exposure, without any distinction between the genotypes. However, when submitted to the cued fear conditioning, ELA2KO displayed lower levels of freezing behavior in the extinction session when compared to WT, but no difference was observed during the conditioning phase. Increased levels of BDNF were found in the prefrontal cortex but not in the hippocampus of ELA2KO mice compared to WT. Finally, in silico analysis indicates that ELA2 is putatively able to cleave BDNF, and incubation of the purified enzyme with BDNF led to the degradation of the later. Our data suggested an anxiogenic‐ and antidepressant-like phenotype of ELA2KO, possibly associated with increased levels of BDNF in the prefrontal cortex.

## Introduction

Elastase-2 (ELA2) is a chymotrypsin-like elastase family member 2A, a serine protease, which is described as an alternative pathway to angiotensin II (ANGII) generation. ELA2 is able to cleave ANGI into ANGII, similarly to angiotensin-converting enzyme (ACE)-mediated reaction. However, ELA2 is also able to generate ANGII directly from angiotensinogen ^1^.

The amino-terminal sequence of the first ten residues of ELA2 is identical to the rat pancreatic elastase-2 (EC 3.4.21.71) and the enzyme was called Ang II-forming ELA2 of rat mesenteric arterial bed ^2^. The cDNAs sequence for pancreatic‐ and rat mesenteric arterial bed-ELA2 showed that they were identical ^3^. ELA2 is considered the only representative of this family of proteases that is secreted outside the digestive tract and is involved in Ang II generation. This enzyme is widely found in several organs such as lung, pancreas, liver, blood vessels (mesenteric and carotid arteries), heart and kidney of rodents ^1,4,5^ and, at lower levels, in the rat prefrontal cortex ^6^). The importance of this functional alternative pathway for Ang II generation was demonstrated in heart, carotid and mesenteric arteries from normotensive and spontaneously hypertensive (SHR) rats ^7^. Also, ELA2 is a key player in ACE-independent dysregulation of the renin-angiotensin system in myocardial infarction and, possibly, other cardiovascular diseases ^8^.

ANGII, acting through its receptors (AGTR1 and AGTR2), can regulate several behavioral responses. For instance, animals lacking AGTR1 receptors, display a facilitated extinction of conditioned fear ^9^. On the other hand, animals lacking AGTR2, did not show any changes in memory tasks, such as the step-down test ^10^, but AGTR2-KOs displayed an anxiogenic phenotype in the elevated plus maze and light-dark box, attenuated by captopril, an inhibitor of ACE ^11,12^. The systemic administration of AGTR1 antagonist losartan exerted biphasic effects in the forced-swimming test, a model predictive of antidepressant-like effect, with lower doses reducing the immobility time ^13^. While captopril was effective in the learned helplessness model ^14^, its effect on the forced-swimming test was contaminated by changes in locomotion ^15^. However, captopril improved the mood of hypertensive patients, comorbidly suffering from depression ^14,16,17^. Moreover, in animal models of anxiety such as the elevated plus maze, the treatment with captopril or losartan exerted an anxiolytic-like effect ^18^. Taken together, these pieces of evidence indicate a complex role for the renin-angiotensin system in the central nervous system.

Thus, the aim of the present study is to describe the phenotype of ELA2KO submitted to models sensitive to anxiety-, depression-like responses, repetitive behaviors, and spatial and cued memory. Based on the behavioral profile of ELA2KO we also analyzed the levels of brain-derived neurotrophic factor (BDNF) in the hippocampus and prefrontal cortex, a neurotrophin that plays a crucial role in these two structures regulating anxiogenic and antidepressant-like responses ^19,20^.

## Experimental procedure

### Animals

adult male and female ELA2KO (38 males/27 females) or C57BL6/j (33 males/33 females) mice, used as wild-type controls (WT), from 12-16 weeks of age at the beginning of the experiments, were employed. The ELA2KO were developed in the Max Delbrück Center for Molecular Medicine, Berlin, Germany; and the colony expanded in the Department of Physiology, at the University of São Paulo in Ribeirão Preto ^21^. The animals were group housed, with free access to food and water, except during experimental sessions. All efforts were made to minimize animal suffering, and the experimental protocols were approved by the local ethics committee at the University of São Paulo in accordance with international guidelines (protocols: 058/2012; 146/2009). Unless otherwise stated, all experiments were conducted in experimentally naive animals, with independent cohorts of male and female mice. However, since no difference was observed between sexes, all behavioral data includes both males and females of each strain.

### Identification of ELA2 mRNA levels in mouse brain

as stated above ELA2 have been described in several tissues in rats and mice. However to our knowledge there is no data pointing to a differential expression in the mouse central nervous system hippocampus or prefrontal cortex. The available literature suggests the expression of *cela2a* gene in rat prefrontal cortex, measured by *microarray* ^6^; or transcriptome sequencing of whole brain ^22^ methods. Thus, samples from prefrontal cortex and hippocampus of 3 males experimentally naive C57BL6/j mice were dissected as described below, washed with PBS and homogenized in Qiazol Lysis ReagentTM (Qiagen, USA). The sample was then incubated with chloroform for 3min at RT. After centrifuged at 15200 x g for 10min at 4oC, the aqueous phase was mixed with isopropanol for 10min at RT and centrifuged at 15200g for 10min at 4^°^C. The pellet was washed with 75% EtOH two time, air dried and dissolved in 20μL MQ-water. Concentration and purity of each RNA sample were determined using NanoDrop (Thermo Scientific). Maxima First Strand cDNA Synthesis Kit for RT-qPCR with dsDNase (#K1672, Thermo Scientific) was used to synthesize cDNA from 2ug of total RNA in the samples. Primers were designed at https://eu.idtdna.com/pages/scitools and purchased from Sigma-Aldrich (Germany).

The following primers for mouse *cela2a* sequence (NM_007919.3) were used for qPCR: forward TCAGGACACTGCTGCTATCT; reverse CAACTACCCTGCTCACATCATC.

For *bdnf exon-9* the following primers were used: forward GAAGGCTGCAGGGGCATAGACAAA; reverse TACACAGGAAGTGTCTATCCTTATG.

Finally, for *beta-actin* (NM_031144) the following primers were used: forward TGTCACCAACTGGGACGATA, reverse: GGGGTGTTGAAGGTCTCAAA.

The PCR method used applied SYBR-Green probe according to manufacturer’s instructions. Briefly, qPCR Master Mix (Thermo Scientific, #K0253) was mixed with 5ul of cDNA and 0.5ul of each primers in Hard-ShellTM 96-well PCR plate (BioRad). The reaction was conducted in duplicates, using thermal cycler (BioRad CFX96 Real-Time System), with initial denaturation at 95^°^C for 10min. Denaturation and amplification were carried out by 45 cycles of 95^°^C for 15s, 63^°^C for 30s and 72^°^C for 30s. ‘No template control’ (NTC) was included to the reaction and melting curve analysis was done. Results were described by cycle threshold (Ct) values.

### Barbering behavior (BB)

the animals were scored for the state of the whiskers and fur, according to a scale for the presence of barbered areas ^23^, based on Garner and colleagues’ ^24^ study of body map for BB. Briefly, an experimenter blind to the genotype scored the animals for the presence or absence of hair loss in any of the following areas: snout, eye area, forehead, chest and neck/back. The presence of barbering in at least one of these areas assigned the analyzed animal to the ‘hair/whisker loss’ group. A total of 52 ELA2KO (26 females) and 54 WT (27 females) at 12-16 weeks of age from our colony were analyzed, all animals were experimentally naive at the time of the analysis and were used in other studies.

### Forced swimming test (FST)

ELA2KO (10 males / 5 females) and WT (7 males / 8 females) were submitted to a 6min session of FST, where the immobility time was measured during the last 4min ^25^. The FST was performed in a cylinder glass (29cm height x 18cm diameter) filled with 10cm water column at 25^°^C. The water was changed between each trial.

### Elevated plus maze (EPM)

ELA2KO (10 males/ 5 females) and WT (7 males/ 8 females) were submitted to EPM for 5min. The percentage of time and entries in the open arm (OAT, OAE respectively) and the entries in the enclosed arm (EAE) were analyzed by ANY-MAZE software (Stoelting, USA). The apparatus used was wood made with two opposite open-arms (30cm x 6cm) and two enclosed arms (30cm x 6cm x 5cm), elevated 50cm above the ground.

### Marble burying test (MBT)

this test was performed in squared cages (28x17x13cm) with 5cm deep sawdust bedding, and 12 dark-green glass marbles evenly distributed. ELA2KO (6 males/5 females) and WT (6 males/6 females) were submitted to a 15min session and the number of buried marbles (more than two thirds) was determined by an experimenter blind to the treatments or genotype ^26^.

### Actimeter test

ELA2KO (7 males/6 females and WT (6 males/6 females) were submitted to a 15min session in the actimeter (Panlab-Harvard Apparatus, Barcelona, Spain). This apparatus consisted in two frames of 32 infrared beams in a transparent square arena to determine the total activity as well as the repetitive/stereotyped movements ^27^.

### Fear conditioning

ELA2KO (5 males) and WT (7 males) were acclimated (2min) to context A (24.8cm long x 25cm wide x 33.5cm tall; black walls with metal grid floor) and submitted to the conditioning protocol by pairing of 5 sound stimulus (30 seconds, 1 Hz, white noise, 80dB) with unconditioned stimulus (1s electrical footshock of 0.6mA). Time trial interval between paired stimuli varied from 20 to 120s. Unconditioned stimulus ended simultaneously with sound. Extinction protocol was conducted in context B (24.8x25x30cm; striped walls and white flat floor) by delivering 21 sound stimuli, the time between stimuli varied from 20 to 60s. The box was cleaned before each trial with a solution of 70% alcohol. Data for both conditioning and extinction protocols were taken as the sum of the time spent in freezing throughout each session and expressed as percentage from WT group. Protocols were performed according to ^28^. Freezing was considered when the animals were immobile, except for respiratory movements. Experiments were recorded and analyzed by an experienced observer blinded to the genotype.

### Sample collection and western-blotting analysis

animals (5 females ELA2KO; 5 females WT) were deeply anesthetized with 4% chloral hydrate injected intraperitoneally, the brains were removed and dissected on ice. Samples from hippocampus (HPC) and prefrontal cortex (PFC) were homogenized in lysis buffer [137mM NaCl; 20mM Tris-HCl; 10% glycerol) containing protease/phosphatase inhibitor cocktail (Sigma-Aldrich, #P2714; #P0044)]. The homogenate was centrifuged (10000 x g) at 4^°^C for 15min and the supernatant collected and stored at −80^°^C. Thirty micrograms of total protein content in each sample was incubated with sample buffer [2% SDS; 5% 2-mercaptoethanol; 10% glycerol; 0.002% bromophenol blue; 125 mM Tris HCl pH= 6.8], separated in acrylamide gel electrophoresis and transferred to a PVDF membrane. Following blockade with 5%BSA in TBST (20mM Tris-HCl; 150mM NaCl; 0.1% Tween20) the membranes were exposed to the primary antibody against BDNF (Millipore, USA, #MABN114) or GAPDH (Santa Cruz, USA, #sc25778) overnight at 4^°^C or HRP-conjugated streptavidin (1:10000, Thermo-Fisher, #21126) for 2h at RT. After washing with TBST the membranes labeled with anti-BDNF or anti-GAPDH, were incubated with secondary HRP-conjugated antibodies for 2h at RT. Following washing the membranes were incubated with 4-chloronaphtol (4CN, Perkin Elmer, #NEL300001EA) for color development or ECL. The dried membranes with 4CN were scanned, and the intensity of bands was determined using ImageJ software (NIH, v.1.49). The chemiluminescence from the membranes exposed to ECL was detected by a CCD camera.

### ELA2 activity on BDNF, *in silico* and *in vitro* analysis

initially we submitted the mature BDNF sequence to PeptideCutter (http://web.expasy.org/peptide_cutter/). However, ELA2 is not one of the currently present options in this server, so we performed the identifications of putative cleavage sites using non-specific chymotrypsin, and later overlapped to the sites with the residues known to be the targets of ELA2 – *i.e.* Leu, Met and Phe. The cleavage at Leu219 residue in BDNF mature sequence would result in an approximately 4kDa lighter chain for this neurotrophin. In order to test this possibility, purified recombinant ELA2 expressed in *E.coli* from mouse sequence (2ug; Biorbyt, UK; #orb246730) was incubated with recombinant human BDNF biotinylated in Lys residues (bBDNF, 200ng, Alomone Labs, Israel, #B-250B) in 100ul of TBS buffer at 37^°^C at 30 or 60min. As control, a sample of bBDNF was incubated with 2ug of BSA in 100ul of TBS for 60min at the same temperature. The samples were than added to sample buffer and submitted to western-blotting for detection of bBDNF by HRP-conjugated streptavidin (as described above).

### Statistical analysis

the incidence of barbering behavior was analyzed by chi-squared test. The FST, EPM, MBT, fear conditioning, actimeter tests, and western-blotting data were analyzed by Student’s t test or Mann-Whitney’s test when necessary. The two-trial test in actimeter was analyzed by two-way ANOVA (with genotype and trial as factors), followed by Fisher’s LSD test. P values below 0.05 were considered statistically significant.

## Results

### Barbering behavior

the chi-squared test indicates a difference on the incidence of barbering behavior in ELA2KO (25 out of 52 scored positive for BB) compared to WT (0 out 54 scored positive): χ^2^= 37.62, p<0.05; as examples can be found in figure 1d. No difference was observed for the incidence of BB in ELA2KO males (12 out 26) and females (16 out 26): χ^2^= 1.24, non-significant, NS.

**Figure 1.**
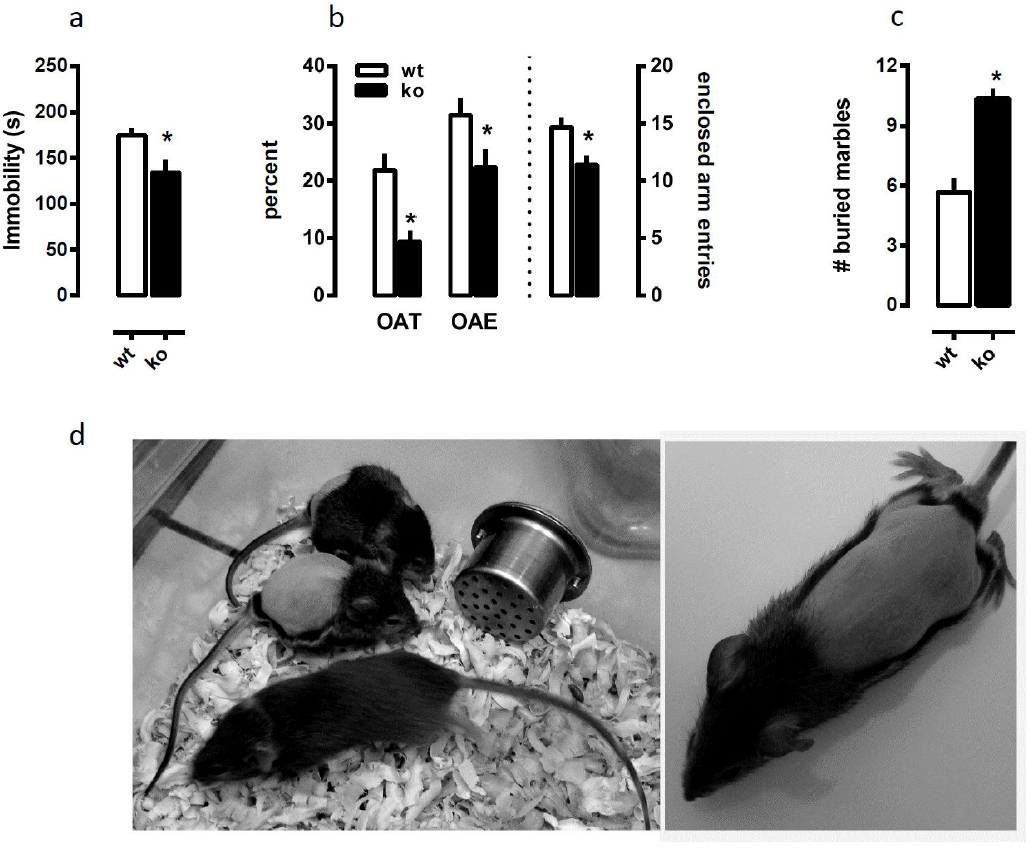
ELA2KO exhibit (a) decreased immobility time in the forced-swimming test (n=15/group); (b) reduced percentages of time spent and entries in the EPM open arm (OAT and OAE respectively) as well as decreased numberx of entries into the enclosed arm (EAE) compared to WT (n=15/group); and (c) increased number of buried marbles in the MBT compared to WT (n=11-12/group). (d) Example of barbering behavior found in ELA2.KO. *p<0.05 from WT.

### Forced swimming test

the Mann-Whitney test indicates a significant effect of genotype, with ELA2KO mice showing decreased immobility time (s) [U=62.00; p= 0.03], as seen in figure 1a.

### Elevated plus maze

as found in figure 1b, the Student’s t test indicates a significant effect of genotype in all the parameters analyzed in EPM; [*i.e.* OAT: t(28)= 3.61; OAE: t(28)= 2.17; EAE: t(28)= 2.84, p<0.05 for all]. Since a significant effect was observed in the number of enclosed arm entries (EAE, considered a locomotion parameter), a covariance analysis of OAT and EAE was performed, and the decreased in OAT in these animals was still significant.

### Marble burying test

the Student’s t test indicates that ELA2KO mice presented an increase in the number of buried marbles [t(21)= 5.52, p<0.05], figure 1c.

### Actimeter test

the Student’s t test indicates a significant effect of genotype on total activity [t(23)= 7.86, p<0.05] and percentage of repetitive movements [t(23)= 3.35, p<0.05], with ELA2KO showing a decrease in these parameters compared to WT, figure 2a-b. The two-way ANOVA of repeated exposure to actimeter indicates a significant effect of time [F(1,30)= 14.93, p<0.05] and genotype [F(1,30)= 69.87, p<0.05], but no interaction between these factors [F(1,30)= 0.13, NS] in the total activity of ELA2KO vs WT, figure 2c.

**Figure 2.**
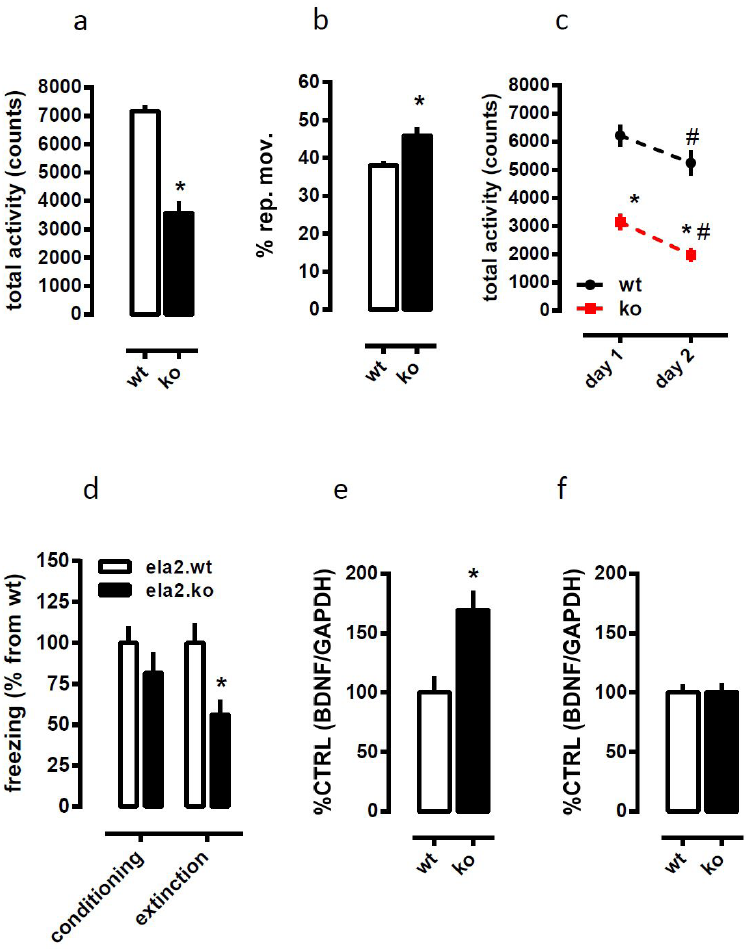
Figure 2. ELA2KO displayed reduced total activity (a) and increased percentage of repetitive movements (b) in the actimeter compared to WT (n=12-13/group). (c) Repeated exposure to the actimeter led to reduced activity in both ELA2KO (red squares) and WT (black circles; n=13-19/group). (d) ELA2KO mice exhibited lower levels of freezing during the extinction phase of contextual fear conditioning (n=5-7/group). ELA2KO exhibited increased levels of BDNF in prefrontal cortex (e), but not in hippocampus (f), compared to WT (n=5/group). *p<0.05 from WT in the same session; #p<0.05 from activity in day 1 in the same genotype.

### Fear conditioning

the Student’s t test indicates a significant difference between the genotype with ELA2KO exhibiting lower levels of freezing in the extinction [t(10)= 2.67, p<0.05] but not in the conditioning phase[t(10)= 1.15, NS] protocols, figure 2d.

### Western blotting data

the Student’s t test indicates a significant increase in the BNDF levels in the prefrontal cortex [t(8)= 3.28, p<0.05] but not in the hippocampus [t(8)= 0.02, NS], figure 2e-f. No changes were observed on the levels of GAPDH in any of the structures analyzed.

### Identification of ELA2 mRNA levels in mouse brain

adult prefrontal cortex and hippocampus display low but detectable levels of *cela2a* mRNA [Mean of Ct values/SD: PFC= 38.88/0.19; HPC= 38.55/0.82; n=3/structure]. As comparison, BDNF Ct values from the same samples were respectively for PFC= 26.56/0.22 and for HPC= 25.66/0.12.

### ELA2 activity on BDNF, *in silico* and *in vitro* analysis

as shown in figure 3a, there is a decrease in the detectable levels of mature BDNF (around 17kDa) following the incubation with ELA2 (30-60min at 37^°^C). The cleavage sites predicted by PeptideCutter, overlapped with those predicted for low-affinity chymotrypsin, are indicated in the figure 3b and the raw data is available in FigShare ^29^.

**Figure 3.**
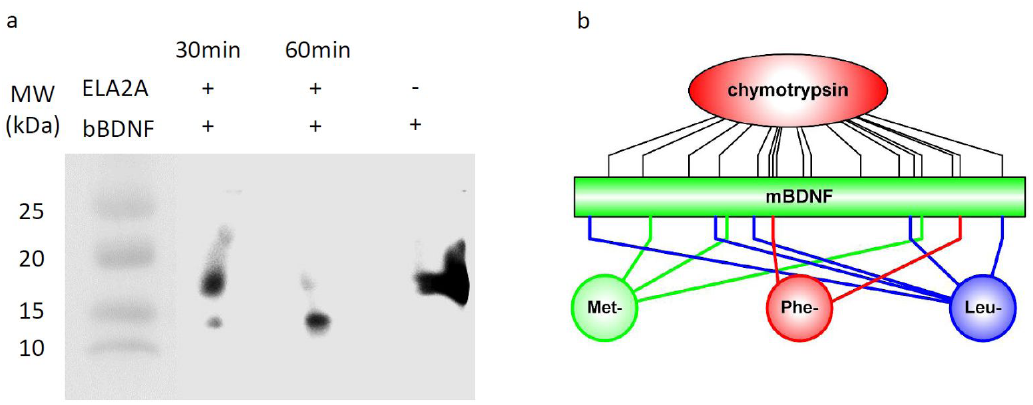
(a) recombinant mouse ELA-2 degrades BDNF. Biotinylated BDNF (BDNF, 200ng) was incubated with purified ELA2 (2ug; 30 or 60min at 37^°^C in TBS buffer). The mixture was resolved by electrophoresis, transferred to PVDF membranes, and developed after incubation with HRP-conjugated streptavidin. As control BDNF (200ng) + 2ug of BSA was kept at 37^°^C for 60min. (b) predicted sites of BDNF cleavage by ELA2, the Leu (blue), Phe (red) or Met (green) residues, were overlapped with low-affinity chymotrypsin cleavage sites.

## Discussion

This is the first study describing that ELA2KO animals exhibit anxiogenic‐ and antidepressive-like phenotype, associated to increased levels of BDNF in prefrontal cortex. These animals, despite no difference in contextual memory or Pavlovian conditioning, do have a more prominent extinction of aversive memories. No difference was observed between males and females in any of the models, thus both sexes were grouped together.

The behavior profile observed in the first experimental approach, the forced-swimming test, indicates an antidepressant-like phenotype. This test was initially designed by Porsolt and colleagues ^30^, and exhibits a high sensitivity to antidepressant compounds (for review see ^31^). However, since it heavily relies on animal’s locomotion, drugs able to increase the general activity, such as stimulants, exert a false positive effect. Therefore, a crucial control for this test is the evaluation of animal’s locomotor behavior. Our data indicates that ELA2KO exhibited decreased activity, evaluated in the actimeter test, compared to WT. Therefore, ELA2KO antidepressant-like phenotype was not biased by increased locomotor activity. Curiously, the treatment with imipramine, a drug able to decrease the immobility time in FST, decreases the expression of *cela2a* gene (GenBank: V01233; SwissProt: P00774) in the prefrontal cortex 2h after a single administration of this antidepressant in rats ^6^.

ELA2KO also exhibited an anxiogenic phenotype, as observed by the reduced time spent and entries in the open arm of the EPM. This test is based on the innate fear that rodents exhibit to open spaces, and have been extensively used to evaluate drugs with putative anxiolytic properties, as well as to the phenotypic characterization of strains and transgenic animals. The analysis of the number of entries in the enclosed arm (EAE) is considered a locomotion parameter in this model (for review of this model see ^32^. Our data showed a decrease in EAE by ELA2KO compared to WT animals. Thus we performed a covariance analysis of the percent time spent in the open arm using the EAE as the covariate (OAT). The effect of genotype on OAT persisted after this analysis, showing that the decrease in time spent in the open arms did not depend on a decrease in general exploratory activity.

In line with the observations in the EPM, ELA2KO exhibited increased burying behavior, and a significant higher incidence of barbering behavior, also suggestive of anxiogenic phenotype. The MBT, similar to EPM, is sensitive to anxiolytics. However, recent studies suggest that it is a not simple measure of anxiety. For instance, the burying behavior observe in MBT is not triggered by a potential threat, as initially interpreted ^26^, but it is a consequence of repetitive behavioral programs. As discussed by Thomas and colleagues ^33^, the nature of the object is not relevant, and there is no correlation between the burying behavior and the parameters observed in the light-dark transition test, a classical model based on the innate fear of rodents to illuminated spaces. Reinforcing the idea that MBT and EPM address different aspects of anxiety traits, we analyzed the repetitive behavior of ELA2KO mice in another test. As found in fig 2, these animals indeed displayed increased repetitive movements in the actimeter. Therefore, it is plausible to interpret that ELA2KO animals also display pro compulsive/repetitive behavior.

The next experimental set was designed to evaluate spatial memory, thus the animals were exposed to the actimeter in two consecutive days. This approach was chosen to minimize the impact of stress and therefore anxiety, as compared for example to Morris’ water-maze or step-down tests. The re-exposure to the actimeter environment led to a decrease in the exploratory behavior of both WT and KO animals, as evidenced by the reduction in the total activity. Since, the reduced activity exhibited by ELA2KO compared to WT persisted in the second trial and concomitantly reduced compared to the first trial, it is plausible to consider that this parameter was not due to the anxiogenic phenotype of ELA2KO. The lack of interaction between the genotype and the trial sessions reinforces the proposition that ELA2KO spatial memory is not compromised. Then, ELA2KO mice were submitted to the cued fear conditioning protocol, where these animals displayed reduced levels of freezing in the extinction session, but no change in the conditioning. Therefore, ELA2KO mice present complex behavioral phenotype related to anxiety, with increased expression of innate fear/anxiety responses, as observed in the EPM, whereas normal or attenuated conditioned fear/anxiety responses and increased extinction of learned fear. We interpret that such phenotype features are consequence of the changes in BDNF levels observed in the prefrontal cortex, but not in hippocampus of ELA2KO.

BDNF and its receptor are considered to play a crucial role in memory, anxiety and depressive-like behavior, this system is suggested to be necessary and sufficient to mediate ^6^. The present study indicates that ELA2KO animals display an anxiogenic‐ and antidepressive-like phenotype, which matches the described phenotype for mice with increased levels of BDNF ^37^. BDNF/TRKB system also plays a core role mediating neuroplastic processes both in hippocampus and cortical areas ^19,38,39^. In line with our findings, mice lacking one of the copies of *bdnf* gene (BDNF+/-), with approximately 50% of BDNF brain levels ^28^; or mice overexpressing the truncated isoform of TRKB ^35^, are more vulnerable to stress, resistant to learning or less flexible to extinguish aversive memories.

As described in the present study, ELA2KO showed increased levels of BDNF in the prefrontal cortex and facilitated extinction in fear conditioning. Some pieces of evidence indicate that BDNF in prefrontal cortex is involved in the expression of learned fear, with distinct roles for its subdivisions. Infusion of BDNF into the infralimbic portion of prefrontal cortex led to a facilitation in cued fear extinction with no effect in the conditioning ^40^. Yet, the selective knockout of BDNF in the prelimbic prefrontal cortex ^41^ also leads to a facilitation in cued fear extinction, without any effects on innate fear responses.

A possible scenario concerns the processing of mature BDNF (mBDNF) by ELA2. It has been described that ELA2 participates in the cleavage of angiotensin I into II ^2^. Based on this evidence, we tested if the proteolytic properties of ELA2 could be applied to mBDNF. First, we submitted the mBDNF sequence to PeptideCutter databank of proteases. Since ELA2 is not among the options in this server, we decided to use low-affinity chymotrypsin as a parameter. As expected, several residues in the mBDNF sequence were suitable to be degraded by the chymotrypsin. Then, we overlapped these residues with putative sites for ELA2 activity, *i.e.* Phe, Leu and Met residues; as indicated in figure 3. Then, we estimated the changes in mBDNF weight starting from the first C-terminal residue. Acting on the C-terminal Leu residues, ELA2 would lead to a 4kDa lighter product from mBDNF. Interestingly, antidepressant-like phenotype and the effect of antidepressant drugs ^19,20,34–36^ we observed a roughly 4kDa lighter product after incubating biotinylated BDNF with purified ELA2, and the blot intensity for this product increases with longer incubation time.

Taken together, the *in silico* data provides an interesting model for the interaction between ELA2 and mBDNF, at least partially confirmed by experimental data, which in turn may provide a mechanism for the increased levels of mBDNF in ELA2KO prefrontal cortex. However, it is important to highlight that to this point we cannot assure that such interaction indeed occur *in vivo,* or have a role, if any, in BDNF metabolism leading to behavioral consequences. Even the literature regarding the existence of *cela2a* gene expression in the central nervous system is scarce. Using transcriptome sequencing (RNA-Seq) analysis of nine tissues from five vertebrates, one study described high levels of *cela2a* expression in the whole rat brain ^22^. Nevertheless, this study did not dissected brain regions putatively implicated in behavioral outcome. Therefore the source of ELA2 cannot be pinned to specific regions or even discard of being from brain blood vessels. In another study, the levels of *cela2a* was observed by *microarray* following treatment with imipramine, in prefrontal cortex of rats ^6^ but no confirmatory analysis by PCR was performed, as well as no comparison with other regions involved in this drug effect such as the hippocampus. In an attempt to replicate and refine the findings in these two studies, we performed PCR analysis of *cela2a* gene expression in two regions of adult mouse brain, *i.e.* the prefrontal cortex and hippocampus, and observed similar levels of expression in both. For comparative reasons we also assessed the levels of *bdnf exon 9* in the same samples. This exon reflects the total levels of *bdnf* gene expression ^42^. We observed approximately 7500-times higher expression of *bdnf exon 9* than *cela2a* in prefrontal cortex and 8500-times in hippocampus. However, to this point, it is still unclear for us the mechanism behind the selective increment of mBDNF protein levels in ELA2KO animals in the prefrontal cortex.

Beyond the changes in BDNF levels, the behavioral phenotype of ELA2KO can be interpreted based on the role of elastases on protease-activated receptors (PAR), mainly PAR-2 ^43^. The function of several G-protein-coupled receptors can be regulated by proteases have been described ^44^. However, specifically to elastases, the literature to such relationship is scarce. Some pieces of evidence point that PAR-2 can be activated by elastases ^45,46^, triggering neurodegenerative effects and increasing neuronal death ^47^. The induction of a cellular insult, such as the microinjection of kainate into hippocampus, leads to infiltration of polymorphonuclear leukocytes (PMNs) in this structure, and the co-culture of neuronal cells with PMNs induces a strong neurotoxic effect, which is attenuated by protease inhibitors ^48^. Yet, this later effect of protease inhibitors only minimally involved serine proteases. As these authors observed, the use of a serine protease inhibitor was not able to counteract the effect of PMNs and kainate insult together in hippocampal cultures, but increased neuronal viability when compared to kainate effect alone.

Interestingly, depressed patients exhibited higher levels of PMN elastase in the serum compared to healthy controls, and such levels are decreased following the treatment with imipramine or moclobemide ^49^. However, there is so far no evidence a such relationship between depression and ELA2. Thus, another curious scenario embeds the ‘leaking’ of serum elastases, putatively including ELA2, to central nervous system following an insult such as hyperactivity of glutamatergic neurotransmission and/or stress.

The increase in glutamate release seems to be a key factor in the responses during, and after stressful events ^50,51^. Moreover, the blockade of NMDA receptors exerts antidepressant-like effect in rats submitted to the forced swimming test ^52^ or the learned helplessness paradigm ^53^. Therefore it is possible to consider the lack of ELA2 could partially prevent the activation of PAR-2, attenuating the deleterious effect of exacerbated glutamatergic neurotransmission.

As pointed before, the main limitation in the present study is that the mechanism how ELA2 contribute to an anxiogenic‐ and antidepressant-like phenotype is not clear. Further studies will be necessary not only to dissect the systems associated to ELA2KO phenotype, but also to understand the interaction between this protease and neuronal development that might underlie such phenotype.

## Acknowledgement

We thank Flávia Salata (USP), Sulo Kolehmainen (UH) and Outi Nikkila (UH) for their technical support, and prof. Eduardo Brandt de Oliveira (Department of Biochemistry, Ribeirão Preto Medical School, USP) for his insightful comments.

